# A Defect in Influenza A Virus Particle Assembly Specific to Primary Human Macrophages

**DOI:** 10.1101/213447

**Authors:** Sukhmani Bedi, Takeshi Noda, Yoshihiro Kawaoka, Akira Ono

**Affiliations:** Department of Microbiology and Immunology, University of Michigan Medical School, Ann Arbor, Michigan, USA; Laboratory of Ultrastructural Virology, Department of Virus Research, Institute for Frontier Life and Medical Sciences, Kyoto University, Kyoto, Japan; Division of Virology, Department of Microbiology and Immunology, Institute of Medical Science, University of Tokyo, Tokyo, Japan; Department of Pathobiological Sciences, University of Wisconsin-Madison, Madison, Wisconsin, USA

**Author notes:** Address correspondence to Akira Ono, Ph.D.

## Abstract

The primary target of Influenza A virus (IAV) is epithelial cells in the respiratory tract. In contrast to epithelial cells, productive infection of most IAV strains is either blocked or highly inefficient in macrophages. The exact nature of the defect in IAV replication in human macrophages remains unknown. In this study, we showed that primary human monocyte-derived macrophages (MDM) are inefficient in IAV release even when compared to a monocytic cell line differentiated to macrophage-like cells, despite comparable levels of expression of viral glycoproteins at the plasma membrane. Correlative fluorescence scanning electron microscopy revealed that formation of budding structures at the cell surface is inefficient in MDM even though clustering of a viral glycoprotein, hemagglutinin (HA), is observed, suggesting that IAV particle assembly is blocked in human MDM. Using an *in situ* proximity ligation assay, we further determined that association between HA and the viral ion channel protein M2 is defective at the plasma membrane of MDM. In contrast, HA and another glycoprotein neuraminidase (NA) associate with each other on the MDM surface efficiently. Notably, the defect in association between HA and M2 in MDM was reversed upon inhibition of actin polymerization by cytochalasin D. Altogether, these results suggest that HA-M2 association on the plasma membrane is a discrete step in the IAV assembly process, which is separable from the association between HA and NA and susceptible to suppression by actin cytoskeleton. Overall, our study revealed the presence of a cell-type-specific mechanism negatively regulating IAV assembly at the plasma membrane.

**Importance:** Identification of host cell determinants promoting or suppressing replication of many viruses has been aided by analyses of host cells that impose inherent blocks on viral replication. In this study, we show that primary human MDM are not permissive to IAV replication due to a defect at the virus particle formation step. This defect is specific to primary human macrophages, since a human monocytic cell line differentiated to macrophage-like cells supports IAV particle formation. We further identified association between two viral transmembrane proteins, HA and M2, on the cell surface as a discrete assembly step, which is defective in MDM. Defective HA-M2 association in MDM is rescued by disruption of the actin cytoskeleton, revealing a previously unknown, negative role for actin polymerization, which is generally thought to play positive roles in IAV assembly. Overall, our study uncovered a host-mediated restriction of association between viral transmembrane components during IAV assembly.

## Introduction

Influenza A virus (IAV) is a negative strand RNA virus that mainly infects and replicates in epithelial cells in the respiratory tract. However, the virus has also been shown to infect other cell types such as macrophages, dendritic cells, and mast cells *ex vivo* (1–3). Host-cell-specific differences have been observed for various properties of IAV including morphology and replication [for example, (4–6)]. These differences could be due to differences in expression levels or functions of host cellular proteins between cell types. In cases where cell-type-specific differences affect productive infection of a virus, detailed comparison between permissive and non-permissive cell types often leads to identification of virus cofactors (5, 7–10) or host factors that restrict replication of viruses (6, 11–14). This approach, which often determines the specific function of the host factor of interest even prior to the identity of the factor, can serve as a complementary approach to genome-wide approaches (15–24).

*Ex vivo* infection studies have shown that in comparison to epithelial cells, murine and human macrophages are less permissive or non-permissive to productive infection of seasonal IAV strains (25–31). In contrast to non-permissive murine macrophages (25, 27, 31, 32), primary human blood-derived or alveolar macrophages support seasonal IAV replication at detectable levels although they are still much less permissive to virus growth than human epithelial cells (26, 28, 29, 32). As for the defective stages of the IAV life cycle, a block at the entry stage of infection has been identified in murine macrophages for most H1N1 strains (25, 27, 31). In addition, the presence of a defect(s) at a later stage has been known for IAV infection in murine macrophages (27, 31). However, there are apparently conflicting data as to whether the defect is at pre- or post-translation stage (27, 31). Moreover, the mechanism in either case has yet to be determined. In contrast to murine macrophages, human macrophages support early stages of replication of all tested IAV strains yet are unable to complete the virus life cycle (31). While the defect appears to be post-translational, the exact nature of this defect in human macrophages and the molecular mechanism behind it are not known.

Determining the nature of the human macrophage-specific defect in IAV replication is likely to advance our understanding of the roles played by cellular functions in late phases of the IAV life cycle and potentially facilitate identification of human host factors involved in this process. In the current study, we used primary human monocyte-derived macrophages (MDM) in order to identify the defective step in IAV replication in human macrophages. We show that MDM support early stages of the IAV assembly process, i.e. trafficking of the viral glycoproteins hemagglutinin (HA), neuraminidase (NA), and the ion channel protein M2 to the plasma membrane, but are inefficient at virus particle formation and subsequent virus release. This defect in virus particle formation and release is specific to primary MDM, since a monocytic cell line THP1 differentiated into macrophage-like cells supports efficient virus particle production. Notably, we observed that the association of HA with M2 on the plasma membrane, as determined by the close proximity of <40 nm, is highly inefficient in MDM relative to the differentiated THP1 cells. In contrast, HA and NA associate efficiently on the surface of MDM. The defective association between HA and M2 is rescued in MDM upon treatment with an actin polymerization inhibitor, cytochalasin D, whereas this defect is recreated in differentiated THP1 cells by treatment with jasplakinolide, which promotes actin polymerization. Overall, this study has identified virus particle formation, more specifically association between HA and M2, as the step defective in IAV life cycle in primary human macrophages and revealed that this host cell-specific block of IAV assembly requires actin polymerization.

## Results

### MDM are inefficient in supporting productive IAV infection relative to differentiated THP1 cells

To determine the extent to which human epithelial cells and macrophages differ in their ability to support productive IAV infection, we compared infectious IAV release from three different human cell types: the lung-derived epithelial cell line A549, the monocytic cell line THP1, which has been differentiated to adopt macrophage-like morphology (dTHP1), and primary monocyte-derived macrophages (MDM). The dTHP1 cells were obtained via treatment of THP1 cells with phorbol 12-myristate 13-acetate (PMA) and vitamin D3 for 2-3 days. A549, dTHP1, and MDM were infected with the laboratory strain A/WSN/1933 (H1N1) (WSN) at MOI 0.01 based on the plaque forming units of virus stocks determined using MDCK cells. At 11 hours post infection (hpi), we observed that virus titers in MDM culture supernatants were up to 100-fold reduced in comparison to that in A549 culture supernatants. Unexpectedly, virus titers in culture supernatants were similar between A549 and dTHP1 cells (Figure 1A). Since dTHP1 cells support influenza virus replication efficiently unlike MDM and yet belong to the same cellular lineage, to facilitate the analyses of MDM-specific defect(s), we chose to compare IAV replication in MDM with that in dTHP1 cells in subsequent experiments. We noticed that while MDM isolated from the vast majority of the tested human donors showed a defect in productive IAV infection relative to dTHP1 at 24 hpi (denoted as Group 1 in Figure 1B), MDM from some donors (denoted as Group 2 in Figure 1B; ~20%) showed no significant difference. Therefore, to identify the MDM-specific defect, the subsequent experiments were performed using MDM from the donors in Group 1. In particular, in the mechanistic experiments (Figures 3–7), we verified in each experiment that MDM used show 10-20 fold reduction in the supernatant virus titers or released vRNA relative to dTHP1 cells at the indicated time point of the corresponding assays (data not shown).

**Figure 1:**
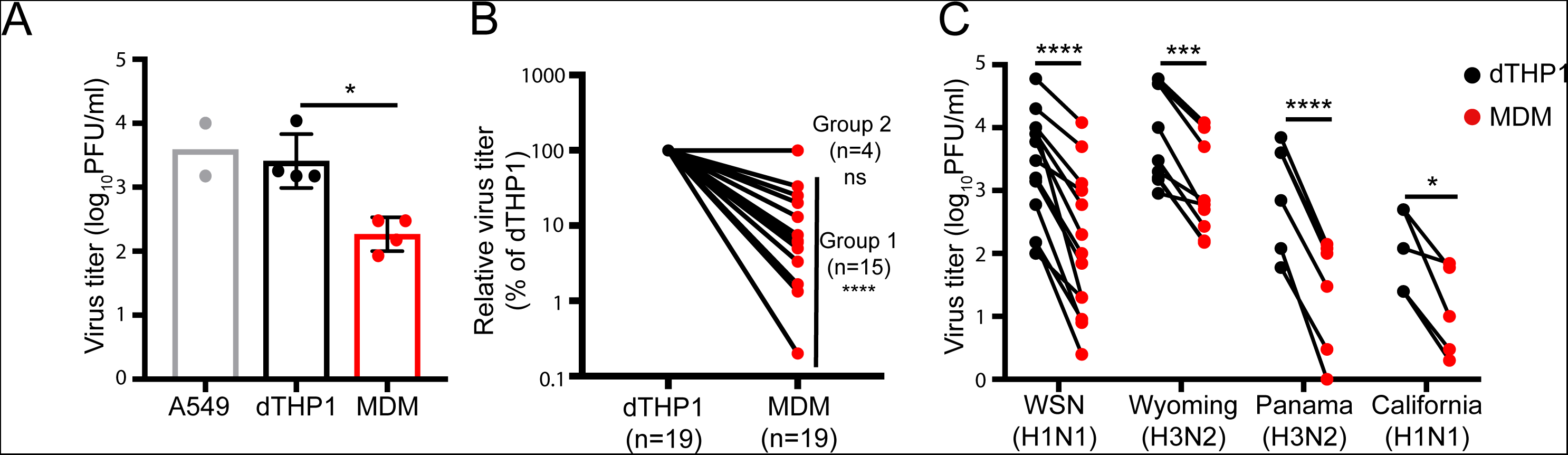
MDM are defective in productive IAV infection. (A) A549 cells, dTHP1 cells and MDM were infected with WSN at MOI 0.01. Infectious virus titers in culture supernatants were measured at 11 hpi. (B) Infectious virus titers in culture supernatants were measured for WSN-infected dTHP1 cells and MDM at 24 hpi. For all tested donors, the relative virus titers in MDM cultures were calculated in comparison to the titer in dTHP1 cell cultures tested in parallel within the same experiment. Two groups of donors (Groups 1 and 2) were denoted based on the reduction in the titers or lack thereof. (C) dTHP1 and MDM (Group 1) were infected with the given IAV strains at MOI 0.01, and infectious virus titers in culture supernatants were measured at 24 hpi. Each circle represents an independently prepared culture. A black and a red circle connected by a line represent each independent experiment. For panel A, data are shown as mean +/- SD. *, P<0.05; ***, P<0.001; ****, P<0.0001; ns, non-significant.

To assess whether other IAV strains also replicate inefficiently in MDM relative to dTHP1 cells, we compared productive infection in dTHP1 cells and MDM of three previously or currently circulating IAV strains, in addition to WSN: A/Wyoming/03/2003 (H3N2) [Wyoming (H3N2)], A/Panama/2007/1999 (H3N2) [Panama (H3N2)], and A/California/04/2009 (H1N1) [California (H1N1)]. Infectious virus titers of all tested IAV strains, as measured by the plaque assay, were reduced by 10-50-fold in MDM in comparison to dTHP1 cells (Figure 1C). These data suggest that MDM are highly inefficient at producing infectious IAV particles in comparison to dTHP1 cells.

### Both efficiency of virus release and infectivity of released particles are impaired in infected MDM relative to infected dTHP1 cells

The results shown above and the results of time-course experiments suggest that infectious virus release is reduced in MDM relative to dTHP1 cells even though flow cytometry using anti-vRNP antibody (clone 61A5 (33)) showed that similar fractions of cells in the cultures are infected (Figure 1 and Supplementary Figure 1). We sought to address whether the reduction in viral titers in MDM culture supernatants is due to a reduction in infectivity of released particles or whether it is due to a reduction in release of physical particles. To this end, we used viral RNA (vRNA) release as a surrogate to measure release of physical particles from dTHP1 cells and MDM. Number of vRNA copies released from MDM were 7-8-fold reduced in comparison to that from dTHP1 cells for all eight vRNA segments. However, there were no significant decrease in cell-associated vRNA levels in MDM relative to dTHP1 cells, indicating that MDM support viral RNA replication and earlier steps as efficiently as dTHP1 cells. vRNA release efficiency was calculated as the ratio of number of vRNA copies in virus pellets from cell culture supernatants to the total number of vRNA copies (cell + virus). For all eight vRNA segments, we observed a 5-10-fold reduction in vRNA release efficiency in MDM relative to dTHP1 cells (Figure 2A). This suggests that efficiency of physical viral particle release from MDM is reduced in comparison to that from dTHP1 cells.

**Figure 2:**
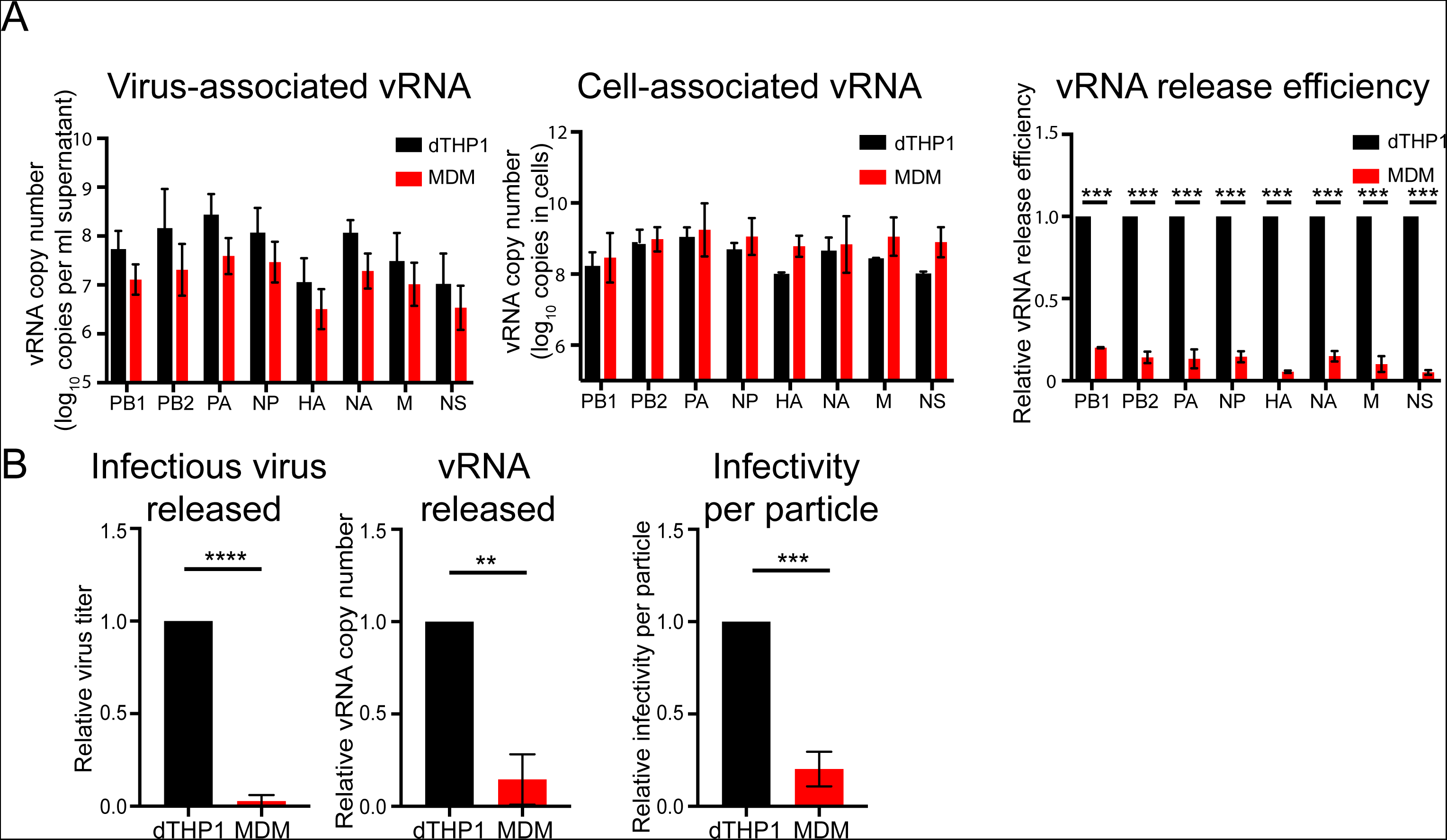
Both efficiency of virus release and infectivity of released particles are reduced in MDM than in dTHP1 cells. dTHP1 cells and MDM were infected with WSN at MOI 0.1. (A) vRNA copy numbers were measured in lysates of virus pelleted from cell culture supernatants and cell lysates at 20 hpi. vRNA release efficiency was calculated as the ratio of number of vRNA copies in virus lysates versus total number of vRNA copies (cell + virus) for each vRNA segment. (B) PB2 vRNA copy number and virus titer were measured in dTHP1 and MDM culture supernatants at 16 hpi. Infectivity per particle was calculated as the ratio of virus titer to PB2 vRNA copy number in culture supernatants. Data are from experiments done with MDM from at least three independent donors and shown as mean +/- S.D. **, P<0.01; ***, P<0.001, ****, P<0.0001.

**Figure 3:**
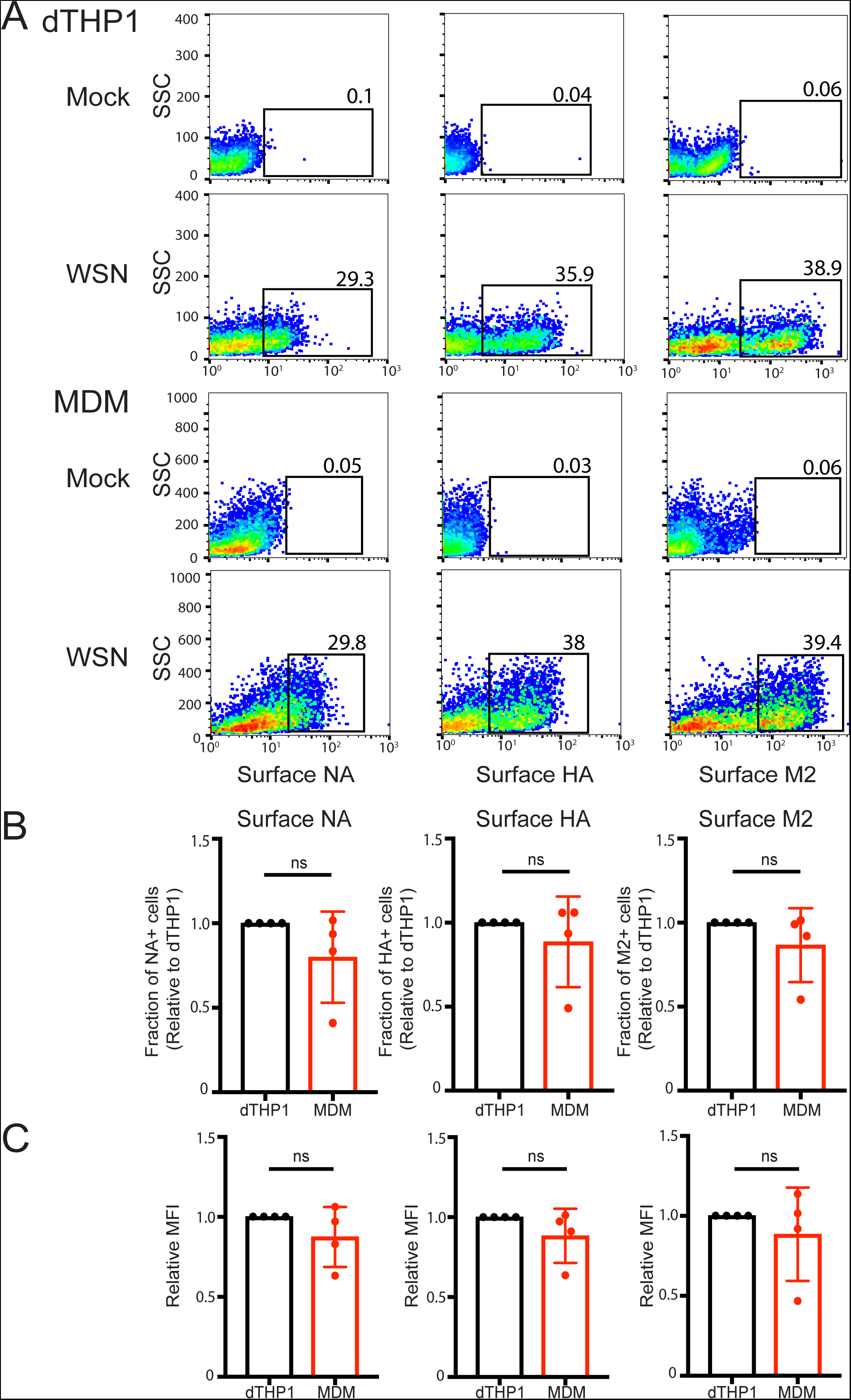
MDM are efficient at trafficking of viral glycoproteins to the cell surface. dTHP1 cells and MDM were infected with WSN at MOI 0.1. (A) Infected cells were analyzed for cell surface expression of HA, NA, and M2 by flow cytometry at 16 hpi. Representative flow plots for mock-infected (top row) and virus-infected (bottom row) cells are shown. Percentages of cells positive for viral proteins (boxed) are shown. Due to differences in the side scatter (SSC) profile between dTHP1 cells and MDM, the Y-axis (SSC) range is different between the two cell types. (B) Percentages of cells positive for surface expression of NA, HA, and M2 are compared between dTHP1 and MDM. (C) Relative MFIs for surface signal of indicated proteins for positive cell populations (gated in panel A) are shown. Data are shown as mean +/- SD and are from at least three independent experiments. ns, non-significant.

Importantly, the reduction in vRNA release (7-8-fold) from MDM does not entirely account for reduction in infectious virus release (up to 50-fold) (Figure 2B). The ratio of released PFU (representing infectious virions) to released vRNA (representing total number of particles) was calculated as the infectivity per particle. Infectivity per particle for virus particles released from MDM was 5-6-fold reduced versus that released from dTHP1 cells (Figure 2B). Overall, our data suggest that the total number of virus particles released as well as the infectivity of released virus particles is reduced in MDM relative to dTHP1 cells.

### Formation of budding structures is inefficient in MDM relative to dTHP1 cells despite similar levels of viral glycoprotein expression at the plasma membrane

We observed that total expression levels of vRNP, HA, and M1 are comparable between dTHP1 cells and MDM (Supplementary Figures 1 and 2), indicating that protein translation and earlier steps are unlikely to be impaired in MDM. Henceforth, we focused on steps post viral protein translation: virus assembly, budding and release. Virus assembly is initiated by targeting of the glycoproteins HA and NA to the plasma membrane (34–36). The third transmembrane protein M2 is also recruited to the assembly sites at the plasma membrane and allows for completion of the virus budding process (35, 37). To determine whether trafficking of the three glycoproteins occurs similarly in dTHP1 cells and MDM, we next compared levels of HA, NA and M2 proteins on the surface of WSN-infected cells. We found that sizes of cell populations positive for surface expression of the three proteins are comparable between MDM and dTHP1 cells (Figures 3A and 3B). The mean fluorescence intensity (MFI) for the three viral proteins in positive cell populations was also similar between MDM and dTHP1 cells (Figure 3C), indicating that trafficking of viral glycoproteins to the plasma membrane is comparable between MDM and dTHP1 cells.

We next asked whether dTHP1 cells and MDM expressing HA on the cell surface support virus particle formation. To address this question in a single cell basis, we performed correlative fluorescence and scanning electron microscopy (CFSEM) in which we first identify cells with surface HA expression using fluorescence microscopy and then examine formation of virus particle-like buds on the surface of the same cells using scanning electron microscopy (SEM). Fluorescence microscopy showed that HA is uniformly distributed on the surface of both dTHP1 cells and MDM with some local accumulation. These HA-enriched clusters or puncta, which likely represent sites of virus assembly, were clearly distinguished on the surface of infected cells after the median filter was applied to the confocal images to remove signal for uniformly distributed non-punctate HA. These HA-enriched sites often corresponded to budding structures with a diameter of approximately 100 nm on the surface of WSN-infected dTHP1 cells (Figure 4A). Very few budding structures with the similar size were observed on the surface of mock-infected cells. MDM also form ~100-nm virus particle-like buds on the surface in HA-positive cells, albeit the number of buds observed in MDM were markedly lower than in dTHP1 cells (Figure 4A). To assess the formation of budding structures quantitatively, we counted the number of HA-positive puncta and the number of virus particle-like buds within the same area (100 µm^2^ in size) of each cell. Even though MDM showed higher numbers of HA-positive puncta on the cell surface than dTHP1 cells, the numbers of virus buds were drastically reduced in MDM relative to dTHP1 cells (Figure 4B). Overall, these results indicate that virus particle assembly/budding are inefficient in MDM despite efficient trafficking of HA, NA and M2 to the plasma membrane.

**Figure 4:**
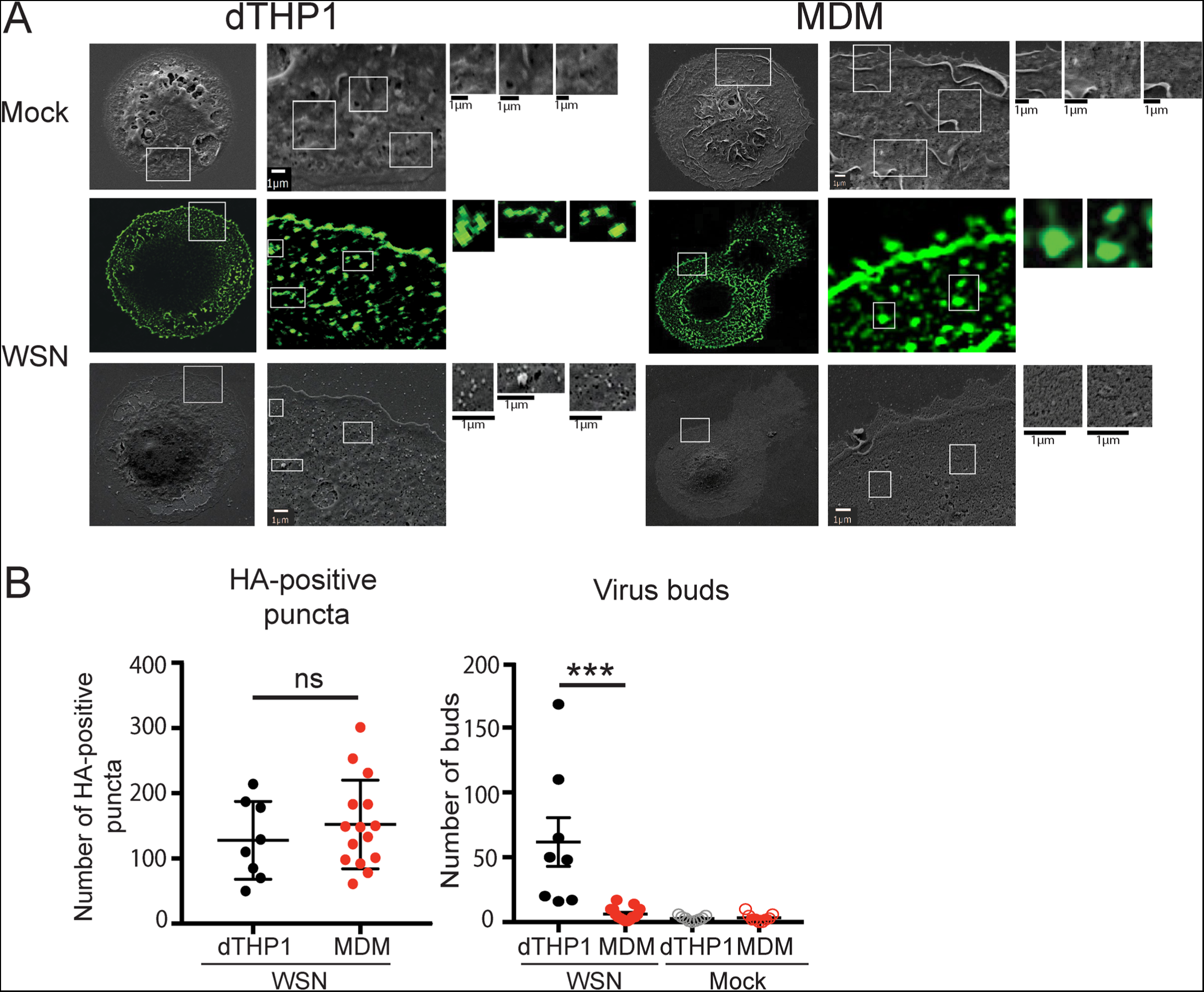
MDM are defective in virus bud formation despite expression of HA on the cell surface. dTHP1 cells and MDM grown on gridded coverslips were infected with WSN at MOI 0.1 for 20 hours. Cells were fixed and immunostained with anti-HA. After identification of HA-positive cells by confocal microscopy, cells were prepared for SEM. The same cells were identified based on grid positions and analyzed by SEM. (A) Representative SEM images for mock-infected and WSN-infected HA-positive cells are shown in the top and bottom rows, respectively. Fluorescence images corresponding to the SEM images of WSN-infected cells are shown in the middle row. Boxed areas are magnified and shown in the right panels. (B) The number of HA-positive puncta identified in fluorescence images (left panel) and ~100-nm buds identified in SEM images (right panel) were counted within the same area (100 µm^2^ in size) in each cell. Data are shown for 8-15 cells from two independent experiments. Error bars represent SEM. ***, P<0.0001; ns, non-significant.

### Association between HA and M2 is impaired in MDM but not in dTHP1 cells

Based on results shown above, we hypothesize that local co-enrichment of HA, NA and M2, which leads to formation of virus assembly sites, are not efficient in MDM relative to dTHP1 cells. To compare formation of the putative assembly sites between dTHP1 cells and MDM, we used *in situ* proximity ligation assay (PLA). PLA allows for detection of two proteins localized within 40-nm distance of each other and has been used to visualize IAV assembly sites on the plasma membrane (38). In addition to measuring PLA signal between the given pair of proteins, we also co-stained cells for cell surface NA to identify infected cells. As a negative control, we performed PLA between HA and transferrin receptor (TfR). TfR does not associate with lipid rafts (39), the plasma membrane microdomains associated with IAV assembly sites (35, 40). Infected dTHP1 cells showed high PLA signal for HA-M2 association. In contrast, infected MDM showed very few PLA spots between HA and M2 (Figures 5A and 5C). As expected, no PLA signal was observed between HA and M2 in mock-infected cells or between HA and TfR in infected cells (Figure 5A). The majority (80-90%) of surface NA-positive MDM and dTHP1 cells express HA and M2 on their surface at comparable levels (Supplementary Figure 3). Therefore, the significant reduction in HA-M2 PLA signal in MDM relative to dTHP1 cells is not due to the lack of expression of HA and/or M2 in NA-positive cells.

**Figure 5:**
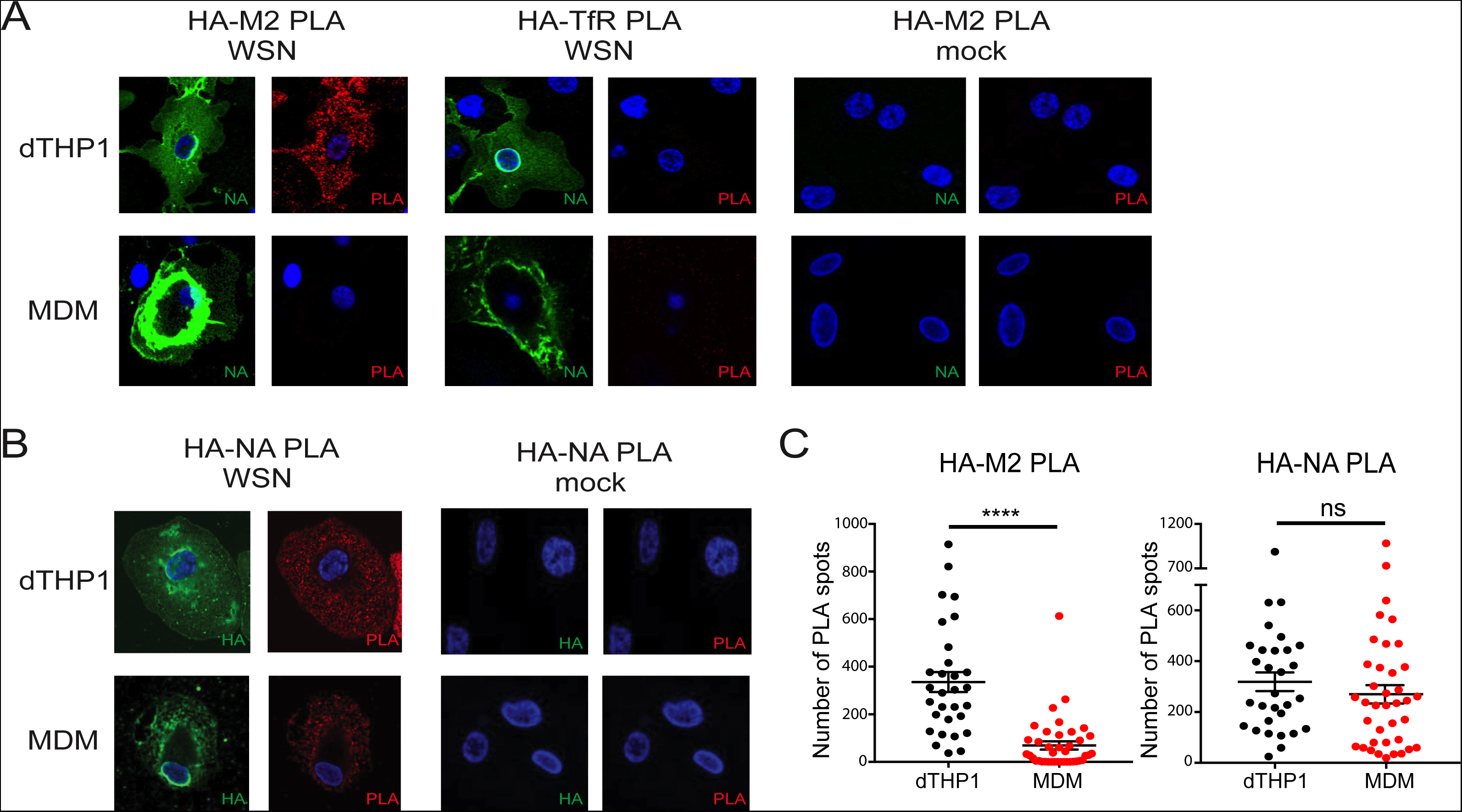
Association between HA and M2 is defective in MDM relative to dTHP1 cells. dTHP1 cells and MDM were infected with WSN at MOI 0.1 for 16 hours. (A) Cells were examined by PLA using goat anti-HA and mouse anti-M2 or goat anti-HA and mouse anti-TfR antibodies. To identify infected cells, surface NA was also detected by rabbit anti-NA. Nuclei were stained with DAPI (blue). Representative maximum intensity projection images, which were reconstructed from z-stacks corresponding to the focal planes ranging from the middle plane of the nucleus to the bottom of the cells, are shown. (B) Cells were examined by PLA using mouse anti-HA and rabbit anti-NA antibodies. Goat anti-HA was used for detection of infected cells. Representative maximum intensity projection images are shown as in panel A. Note regions of intense NA (in A) and HA (in B) signal on the surface of MDM due to the presence of membrane ruffles. (C) Number of PLA spots were counted for each cell. Data are shown for three independent experiments, and 8-10 cells were analyzed per experiment. These experiments were performed in parallel with the experiments shown in Figure 3 using MDM from the same donors. Error bars represent SEM. *, p<0.05; ****, p <0.0001; ns, non-significant.

To determine whether the defect in association between transmembrane proteins in MDM is specific to HA and M2 or whether association between other pairs of viral transmembrane proteins is defective as well, we next measured PLA signal between HA and NA. In this case, to identify infected cells, we co-stained cells for cell surface HA using an antibody different from the one used for PLA. PLA signal between HA and NA was similar for dTHP1 cells and MDM, suggesting that HA and NA associate with each other as efficiently on the surface of MDM as on dTHP1 cells (Figures 5B and 5C). Overall, our data indicate that association between HA and M2 is a virus assembly step specifically impaired in MDM.

### Inhibition of actin polymerization restores HA-M2 PLA in MDM, while promotion of actin polymerization reduces HA-M2 PLA in dTHP1 cells

HA associates with lipid rafts on the plasma membrane, while M2 mainly localizes in non-lipid raft areas (35, 40). It is suggested that M2 is recruited to cholesterol-rich lipid rafts during IAV particle assembly (41, 42); however, host cell functions and factors that regulate this step are not known. It is possible that in MDM, HA-containing plasma membrane microdomains stay segregated from those containing M2, leading to defective association between the two glycoproteins. The cortical actin cytoskeleton, a network of filaments that underlies and interacts with the plasma membrane, is suggested to play a role in formation and maintenance of plasma membrane microdomains (43–45). Therefore, we next asked whether the actin network regulates the association between HA and M2 in dTHP1 cells and MDM. Infected cells were treated with cytochalasin D (Cyto D), an inhibitor of actin polymerization, at 14 hpi for 2 hours, fixed, and examined for HA-M2 association using PLA. Infected dTHP1 cells showed high PLA signal for HA-M2 association under both untreated and Cyto D-treated conditions. As observed in Figure 5B, untreated MDM showed very few PLA spots between HA and M2. In contrast, Cyto D-treated MDM showed PLA signal between HA and M2 at levels similar to that observed for dTHP1 cells (Figures 6A and 6B). No PLA signal was observed between HA and TfR in untreated or Cyto D-treated dTHP1 cells and MDM. The increase in HA-M2 PLA signal upon Cyto D treatment of MDM was not due to an increase in surface expression of HA and M2 in drug-treated cells, as shown by the flow cytometry analysis (Supplementary Figure 4).

**Figure 6:**
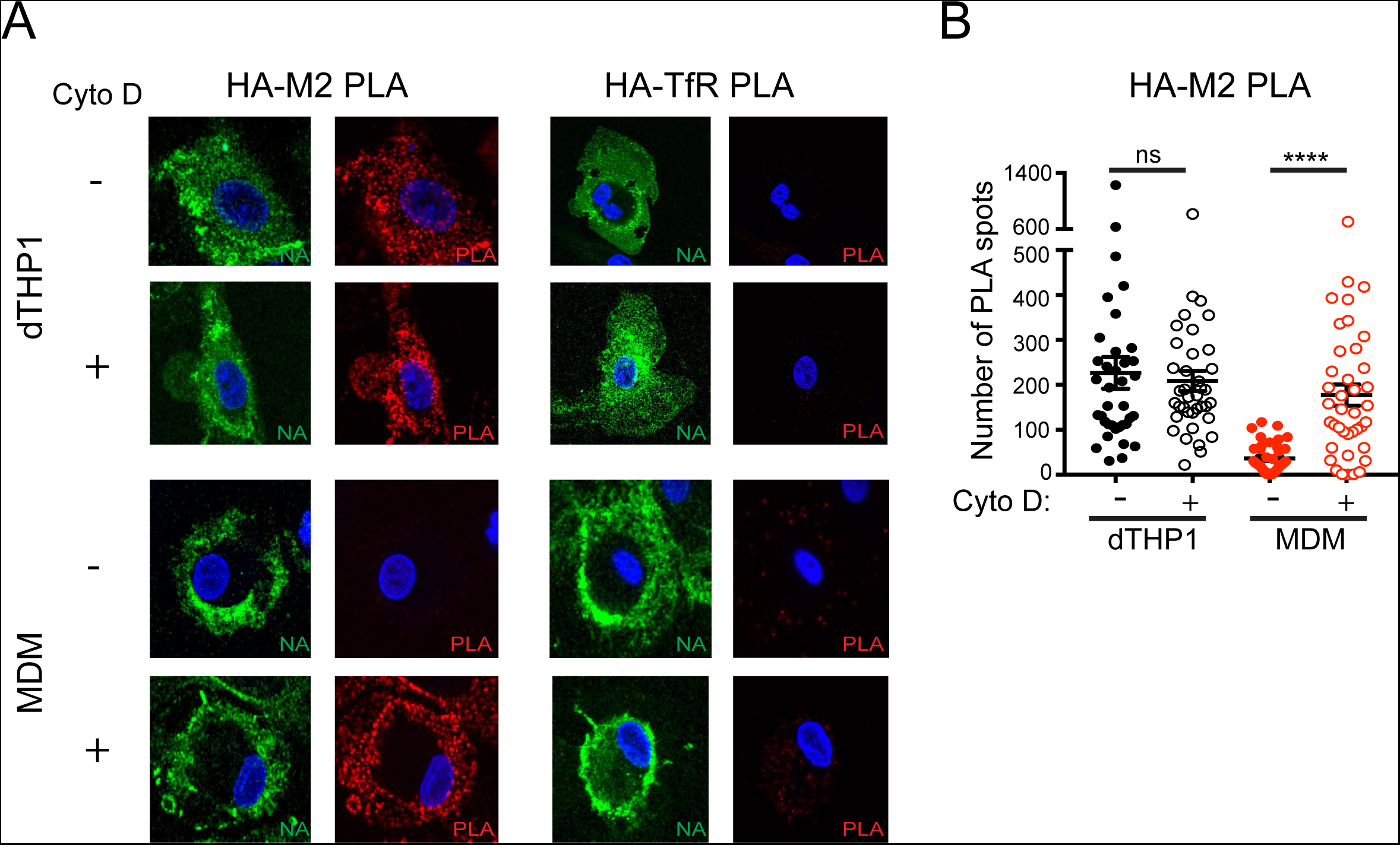
Cytochalasin D treatment restores HA-M2 PLA in MDM to levels comparable to that in dTHP1 cells. dTHP1 cells and MDM were infected with WSN at MOI 0.1. At 14 hpi, cells were treated with vehicle control (DMSO) or 20 µM cytochalasin D (Cyto D) for 2 hours before fixation. (A) Cells were analyzed as in Figure 5A. Representative maximum intensity projection images are shown. (B) Number of PLA spots were counted for each cell. These experiments were performed in parallel with the experiments shown in Supplementary Figure 4 using MDMs from the same donors. Data are from at least three independent experiments, and 8-10 cells were analyzed per experiment. Error bars represent SEM. ****, p <0.0001; ns, non-significant.

Since inhibition of actin polymerization restores association of HA and M2 in MDM, we next asked whether promoting actin polymerization inhibits HA-M2 association in dTHP1 cells. To this end, infected dTHP1 cells were treated with jasplakinolide (Jasp), which nucleates and stabilizes actin polymerization, at 14 hpi for 2 hours and examined for HA-M2 association using PLA at 16 hpi as in Figure 6. Two hours of Jasp treatment reduced HA-M2 PLA in 50% of the examined dTHP1 cells, while the remaining infected cell population showed HA-M2 PLA signal comparable to that in untreated cells (data not shown). We reasoned that high HA-M2 PLA signal in 50% of Jasp-treated dTHP1 cells is due to pre-existing association between HA and M2 at the time of Jasp addition. Therefore, we next examined the effect of Jasp on HA-M2 association at an earlier time point in infection when pre-existing HA-M2 coclusters are unlikely to be abundant. We treated infected dTHP1 cells with Jasp or Cyto D at 10 hpi for 4 hours and examined for HA-M2 association using PLA at 14 hpi. Under these conditions, most Jasp-treated dTHP1 cells showed reduced HA-M2 PLA signal, in comparison to untreated or Cyto D-treated cells (Figures 7A and 7B). Since the defect in HA-M2 association is rescued upon inhibition of actin polymerization in MDM while it is induced upon stabilization of actin in dTHP1 cells, our data overall highlight a role for actin polymerization in suppressing association between HA and M2 at the plasma membrane.

**Figure 7:**
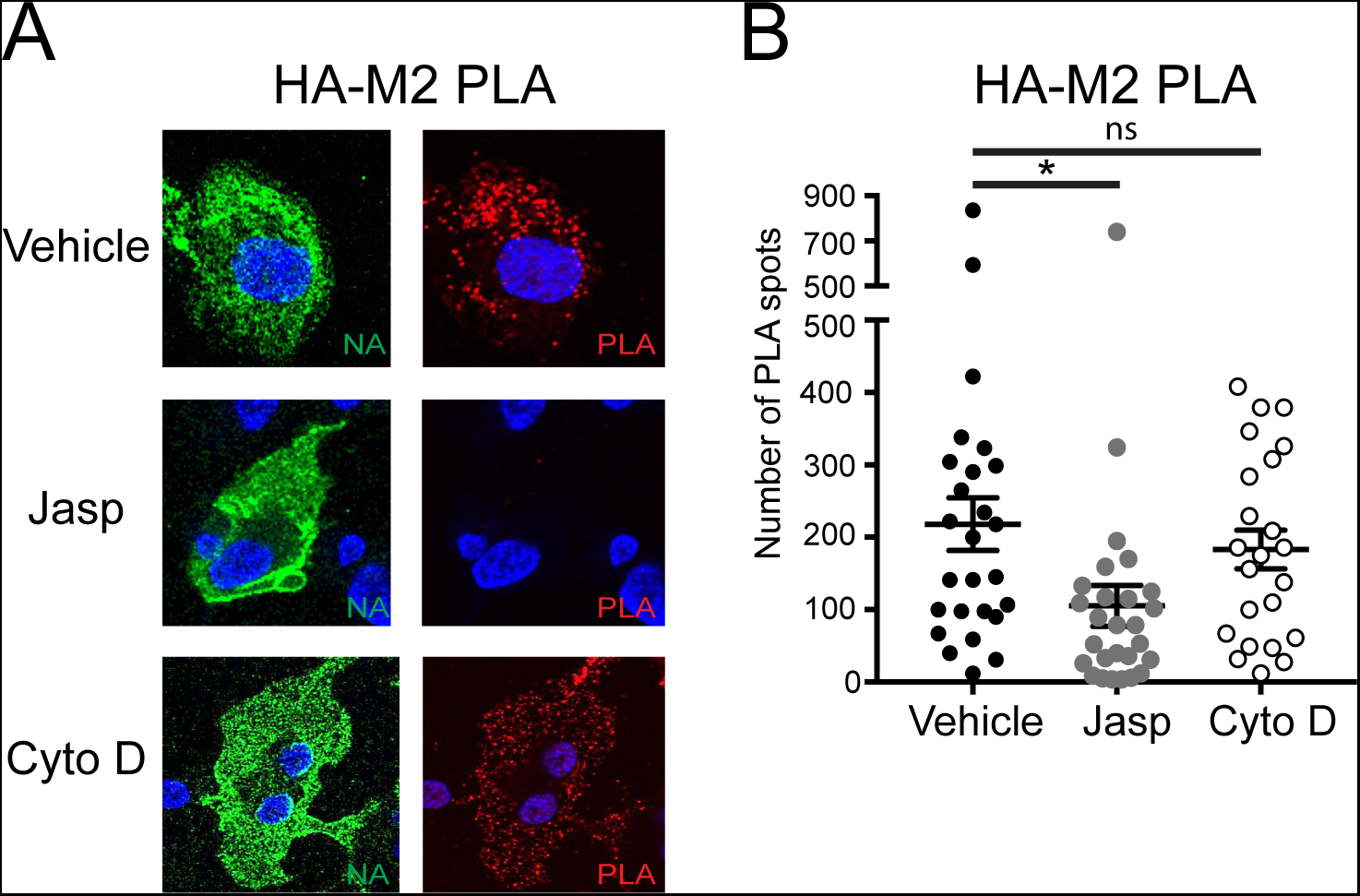
Jasplakinolide treatment reduces HA-M2 PLA in dTHP1 cells. dTHP1 cells were infected with WSN at MOI 0.1. At 10 hpi, cells were treated with vehicle control (100% Ethanol), 1 µM Jasplakinolide (Jasp), or 20 µM Cyto D for 4 hours before fixation. (A) Representative maximum intensity projection images are shown as in Figure 5A. (B) Number of PLA spots were counted for each cell. Data are from at least three independent experiments, and 7-10 cells were analyzed per experiment. Error bars represent SEM. *, p <0.05; ns, non-significant.

## Discussion

In a previous study, a post-translational defect in productive IAV infection was observed in human MDM (31). A similar defect was also reported for murine macrophages in one study (27) but not the other (31). The exact nature of these defects that lead to inefficient IAV release has not been determined. Here, we have shown that despite efficient trafficking of the viral glycoproteins to the cell surface (Figure 3), infectious virus particle formation at the plasma membrane is inefficient in human MDM (Figures 2 and 4). The current study further identified HA-M2 association as an IAV assembly step suppressed in MDM (Figure 5). This restriction is specific to primary macrophages, as the THP1 monocytic cell line differentiated to macrophage-like cells (dTHP1 cells) support HA-M2 association and efficient IAV production. Notably, defective HA-M2 association in MDM can be reversed by the disruption of actin polymerization, suggesting a role for the actin cytoskeleton in suppressing this step of IAV assembly in MDM (Figure 6). Consistent with the restrictive role of actin polymerization, the HA-M2 association in dTHP1 cells was blocked upon a treatment that promotes actin polymerization (Figure 7).

Previous studies observed strain-specific differences for IAV replication in human macrophages. Some strains such as highly pathogenic H5N1 and pandemic 1918 strains can replicate in macrophages albeit at a lower efficiency than in epithelial cells (25, 26, 29, 32). Marvin *et al.* recently reported that the laboratory strain WSN is able to overcome blocks in IAV replication in human macrophages, while replication of the A/California/04/2009 strain is completely blocked (31). In our study, we observed that all tested strains, including WSN and A/California/04/2009, released significantly lower titers in MDM than in dTHP1 cells (Figure 1C). Thus, it is likely that the cell-type-specific difference observed in this study is distinct from the previously reported strain-specific difference.

In addition to identifying the defective step for IAV replication in primary human macrophages, our study also lends mechanistic insights into the assembly and budding process of IAV in host cells. IAV is thought to assemble in cholesterol-enriched microdomains, or membrane rafts, of the plasma membrane in host cells (46–49). HA and NA accumulate at these assembly sites (34–36, 40, 50), also known as the budozones, while the third transmembrane protein M2 is suggested to localize at the edge of the budozone (37, 42, 51). Coclustering between HA and M2 has been observed at steady state in epithelial cells (41, 51, 52). However, the sequence of events leading to recruitment of M2 to the budozone is unknown. Whether there is a mechanism regulating these events, other than simple diffusion over the plasma membrane, also remains to be determined. Our PLA data suggest that association between HA and NA likely precedes association between HA and M2. Consistent with this possibility, a recent study showed that NA but not M2 accelerates HA trafficking to the apical surface of epithelial cells, presumably through co-trafficking (50). Thus, recruitment of M2 to assembly sites enriched in HA (and perhaps NA) is a discrete and host-cell-dependent step in the IAV assembly process.

The actin cytoskeleton has been implicated in assembly of IAV particles, in particular formation of filamentous particles (4, 53, 54). However, the actin-dependent mechanism(s) regulating IAV assembly are not well understood. Previous studies have shown that disruption of actin dynamics by both inhibition of actin polymerization (4, 53) (by latrunculin A or cytochalasin D) and promotion of actin polymerization (53) (by jasplakinolide) disrupts filamentous IAV assembly. In contrast, our study showed that drugs inhibiting actin polymerization and depolymerization have distinct and opposing effects on HA-M2 association: blocking actin polymerization in IAV non-permissive cells (MDM) restores HA-M2 association, whereas enhancement of polymerization in permissive cells (dTHP1 cells) reduces HA-M2 association. Therefore, it is likely that distinct actin-dependent mechanisms regulate the association between HA and M2 at the plasma membrane and formation of filamentous particles. The regulation of HA-M2 association may depend on the segregation of HA- and M2-enriched plasma membrane microdomains by the actin cytoskeleton. Consistent with this possibility, previous studies support a role for actin polymerization in maintaining HA-enriched microdomains compact and dense (55, 56).

The cytoplasmic domain of M2 comprises of an amphipathic helix, which plays a role in scission of the IAV particle after budding (37, 51). Previous studies have described the role of M2 in assembly and scission of virus buds (51, 57), vRNP incorporation (58–61), and M1 binding (41, 58, 61–63) using virus strains that either lack M2 expression or express M2 cytoplasmic tail mutants. Interestingly, in epithelial cells, virus lacking M2 or expressing mutant M2 proteins is still able to initiate particle assembly and budding, presumably mediated by HA and NA (34, 64). However, these buds adopt an abnormal morphology and/or fail to undergo scission or release (37, 51, 60), latter of which results in accumulation of particles at the cell surface. In contrast, MDM showed very few buds on their surface (Figure 4), suggesting that in MDM, an earlier role for M2 in virus particle assembly is more prominent than in epithelial cells. Consistent with this early role of M2 in IAV assembly, M2 functions in recruitment of M1 and vRNP to assembly sites (41, 58–63), which is important for initiation of IAV particle assembly or elongation of filamentous particles (58, 60, 65–71). A defect in incorporation of M1 and/or vRNP into budding virus particles due to the failure of M2 recruitment may also explain the reduction in infectivity per particle observed for MDM-derived virus relative to dTHP1-derived virus (Figure 2B). We also do not rule out the possibility that a failure in M1 association with the assembly sites may contribute to the observed defect in M2-HA association specific to MDM. However, we note that there was no major difference in distribution of M1 relative to the site of HA accumulation at the plasma membrane between MDM and dTHP1 cells (unpublished observations).

Overall, in this study, we have compared IAV replication in MDM with that in dTHP1 cells and found that MDM replicate vRNA, express viral proteins, and traffic HA, NA and M2 to the plasma membrane at levels similar to those in dTHP1 cells. However, MDM are defective in assembling virus particles, likely due to actin-dependent suppression of association between the viral transmembrane proteins HA and M2. Comparison of actin regulatory mechanisms operating in MDM and dTHP1 cells, which are of the same lineage, will likely facilitate identification of additional host cellular factors involved in the assembly stage of the IAV life cycle.

## Materials and Methods

### Cells and reagents

Monocytes were isolated by plate adhesion from peripheral blood mononuclear cells, which were obtained from buffy coats derived from unidentified healthy donors (New York Blood Center, NY). Cells were cultured in RPMI 1640(Gibco) supplemented with 10% fetal bovine serum (FBS, Hyclone) for 7 days before they were used for experiments. THP1 (ATCC® TIB202™) cells were cultured in RPMI 1640 supplemented with 10% FBS, 1 mM Sodium Pyruvate (Gibco) and 0.05 mM 2-mercaptoethanol. To generate differentiated THP1 cells (dTHP1), THP1 cells were cultured in the medium containing 0.1 µM phorbol 12-myristate 13-acetate (PMA; Sigma) and 0.1 µM Vitamin D3 (Sigma) for 2-3 days. Madin-Darby canine kidney (MDCK) cells were provided by Dr. Arnold S. Monto (University of Michigan) and were cultured in DMEM (Gibco) supplemented with 10% FBS and 25 mM HEPES. Human lung carcinoma cell line A549 was provided by Dr. Mike Bachman (University of Michigan) and was cultured in DMEM (Gibco) supplemented with 10% FBS and 25 mM HEPES. Human embryonic kidney-derived 293T cell line (ATCC) was cultured and maintained in DMEM (Lonza) supplemented with 10% FBS.

The following antibodies were used for immunofluorescence microscopy: mouse anti-HA monoclonal antibody (clone C179 (72); Takara), mouse anti-M2 monoclonal antibody (clone 14C2 (73); Thermofisher), mouse anti-vRNP monoclonal antibody (clone 61A5 (33); a kind gift from Dr. Fumitaka Momose, Kitasato University), goat anti-HA antiserum (BEI NR-3148), mouse anti-transferrin receptor (TfR) monoclonal antibody (clone M-A712; BD Biosciences). Rabbit anti-NA antiserum was a kind gift from Dr. Christopher Brooke (University of Illinois). All secondary antibodies used for immunofluorescence were purchased from Thermofisher. Cytochalasin D and Jasplakinolide were purchased from Sigma and re-constituted in DMSO and 100% Ethanol, respectively.

### Plasmids and virus stocks

A/WSN/1933 (H1N1) virus was generated by reverse genetics (74) using the 8 pPolI plasmids encoding different segments of IAV genome and the 4 pCAGGS plasmids that express the PA, PB1, PB2, and NP proteins. The titers of the stocks were determined using the plaque assay with MDCK cells. A/Wyoming/03/2003 (H3N2) [Wyoming (H3N2)], A/Panama/2007/1999 (H3N2) [Panama (H3N2)], and A/California/04/2009 (H1N1) [California (H1N1)] viruses were kind gifts from Dr. Arnold S. Monto (University of Michigan) and were received as low passage stocks (less than 5 passages in MDCK cells) of virus isolated from clinical specimen. Virus infection was performed and monitored using the plaque assay and flow cytometry as described in Supplementary Information.

### Measurement of vRNA levels

Virus-containing cell culture supernatants were centrifuged at 3000 rpm for 5 minutes in a microfuge, filtered through a 0.45-µm filter, and subjected to ultracentrifugation at 30,800 rpm (AH650 swinging bucket rotor, Thermofisher) for 90 minutes to prepare virus pellets. Virus and cell-associated vRNA was measured using a previously described protocol (75). Briefly, total RNA was extracted from virus pellets and cell lysates using TRIzol reagent (Ambion) according to the manufacturer’s protocol. Complementary DNA was generated using random hexamer priming and the SuperScript III First-Strand Synthesis System (Invitrogen). Quantitative PCR was performed on a CFX96 Real Time PCR system (Biorad) using Platinum SYBR Green pPCR SuperMiX-UDG (Thermo Scientific Fisher). Serial ten-fold dilutions of pPolI plasmids containing specific viral gene of WSN were used to generate a standard curve for quantification of cDNA copy number based on cycle threshold (Ct) values. The primer sequences are shown in Supplementary Information.

### Correlative fluorescence and scanning electron microscopy (CFSEM)

CFSEM experiments were performed as described before (76). Briefly, cells cultured on gridded coverslips (Bellco Biotechnology) were infected with WSN at MOI 0.1. Cells were fixed with 4% PFA in PBS at 20 hpi. After rinsing in PBS, quenching of PFA with PBS containing 0.1 M glycine (Sigma), and blocking with PBS containing 3% bovine serum albumin (BSA, Sigma), cells were immunostained with mouse anti-HA and fluorescently labeled secondary antibody. Cells were imaged using a Leica Inverted SP5X Confocal Microscope with a 40× PL APO objective and 10-20× scanning zoom. After fluorescence imaging, cells were fixed with PBS containing 2.5% glutaraldehyde (Electron Microscopy Sciences), stained with 1% OsO_4_, dehydrated in a series of ethanol washes, rinsed in hexamethyldisilazane (Electron Microscopy Sciences) and allowed to dry overnight. Coverslips were affixed to specimen mounts and sputter coated with gold for 90 s (Polaron). Cells were identified by their location on the gridded coverslip and imaged on an Amray 1910FEG scanning electron microscope at 5-10 kV. Fluorescence and SEM images were roughly brought into registration by scaling and rotating images in Adobe Illustrator, similarly to other correlative fluorescence/SEM studies (76). Landmarks used for registration included cell edges. Cell surface structures visible in SEM were manually classified as virus-like buds if they appeared spherical and near 100 nm in diameter. To identify HA clusters in fluorescence images unambiguously, we removed uniform non-puncta HA signal from the images. To do this, we calculated a 20-pixel radius median filter and subtracted the median filtered image from the original using the *Image Calculator* function in ImageJ. Number of HA-positive puncta was measured in the background-subtracted fluorescence images using the *Analyze particle* function in ImageJ. Since MDM have substantial membrane folds on the cell surface especially towards the center of the cell, we focused on areas towards the edge of the cells, which have a flatter topology, for quantification of efficiency of virus bud formation.

### *In situ* Proximity Ligation Assay (PLA)

PLA was performed using Duolink® PLA fluorescence kit following the manufacturer’s instruction (Sigma). Cells fixed with 4% PFA (non-permeabilized) were incubated with following primary antibody combinations: goat anti-HA and mouse anti-M2 for PLA and rabbit anti-NA for identification of infected cells; mouse anti-HA and rabbit anti-NA for PLA and goat anti-HA for identification of infected cells; or goat anti-HA and mouse anti-TfR for PLA and rabbit anti-NA for identification of infected cells. Detection of PLA signals and identification of infected cells were performed using PLA probes specific to goat, mouse or rabbit IgG and AlexaFluor-488-labeled secondary antibody recognizing anti-NA or anti-HA, respectively. Cells were observed using a Leica Inverted SP5X Confocal Microscope System with a 63X objective. Z-stacks extending from the focal plane corresponding to the middle plane of the nucleus (identified by DAPI staining) to the bottom of cells were acquired for each cell, and the maximum intensity projection for each cell was constructed using ImageJ. The PLA signal in projection images was thresholded to eliminate weak and hazy background signal in the nucleus, and the number of PLA-positive spots was counted using the *Analyze particle* function in ImageJ.

### Statistical analysis

Statistical analyses were performed using GraphPad Prism version 7. Two-tailed paired student *t* test was used to calculate p-values in Figures 1–3 and supplementary Figures 1–4. Two-tailed unpaired student *t* test was performed in Figures 4–7.

## Acknowledgments

We would like to thank the members of our laboratories for helpful discussions and critical review of the manuscript. We would like to thank Drs. Bachman, Brooke, Momose, and Monto for reagents. The following reagent was obtained through BEI Resources, NIAID, NIH: Polyclonal Anti-Influenza Virus H1 (H0) Hemagglutinin (HA), A/Puerto Rico/8/1934 (H1N1), (antiserum, Goat), NR-3148. This work was supported by funding from the National Institutes of Health (R01 AI071727) (to A.O), the Takeda Science Foundation (to T.N), and Leading Advanced Projects for medical innovation (LEAP) from the Japan Agency for Medical Research and Development (AMED) (JP17am0001007) (to Y.K.). S.B. was supported by a Clayton Willison- and Emma Elizabeth Willison-Endowed Graduate Fellowship.

## Supplementary Information

**Supplementary Figure 1:**
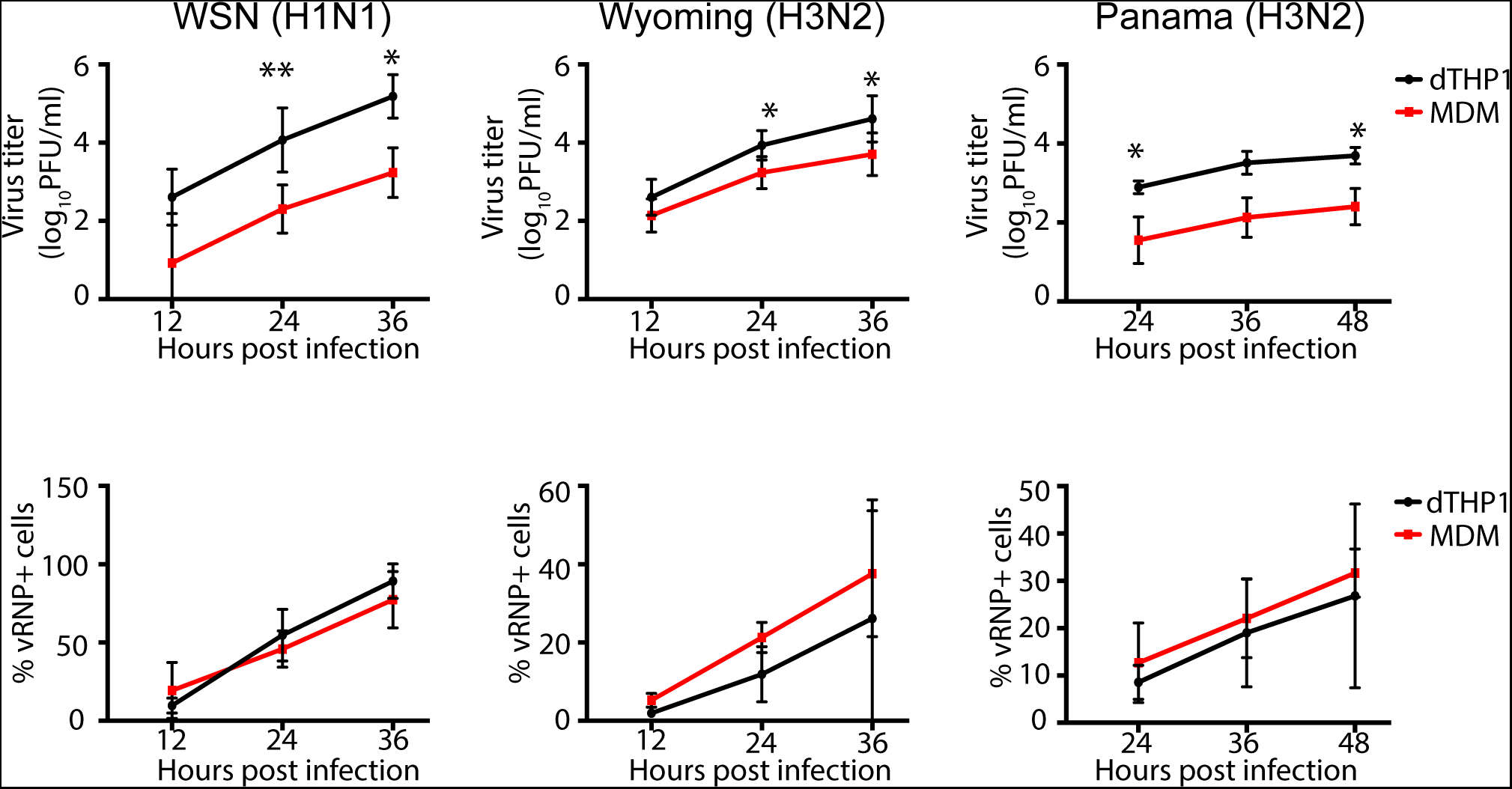
Infectious virus release is reduced in MDM cultures relative to dTHP1 cells despite similar number of infected cells in both cultures. dTHP1 cells and MDM were infected with given IAV strains at MOI 0.01. At the indicated time points post infection, culture supernatants and cells were collected and assessed for virus titer (top panels) and % vRNP+ cells (lower panels), respectively. Data from at least three independent experiments are shown as mean +/- SD. *, P<0.05; **, P<0.01.

**Supplementary Figure 2:**
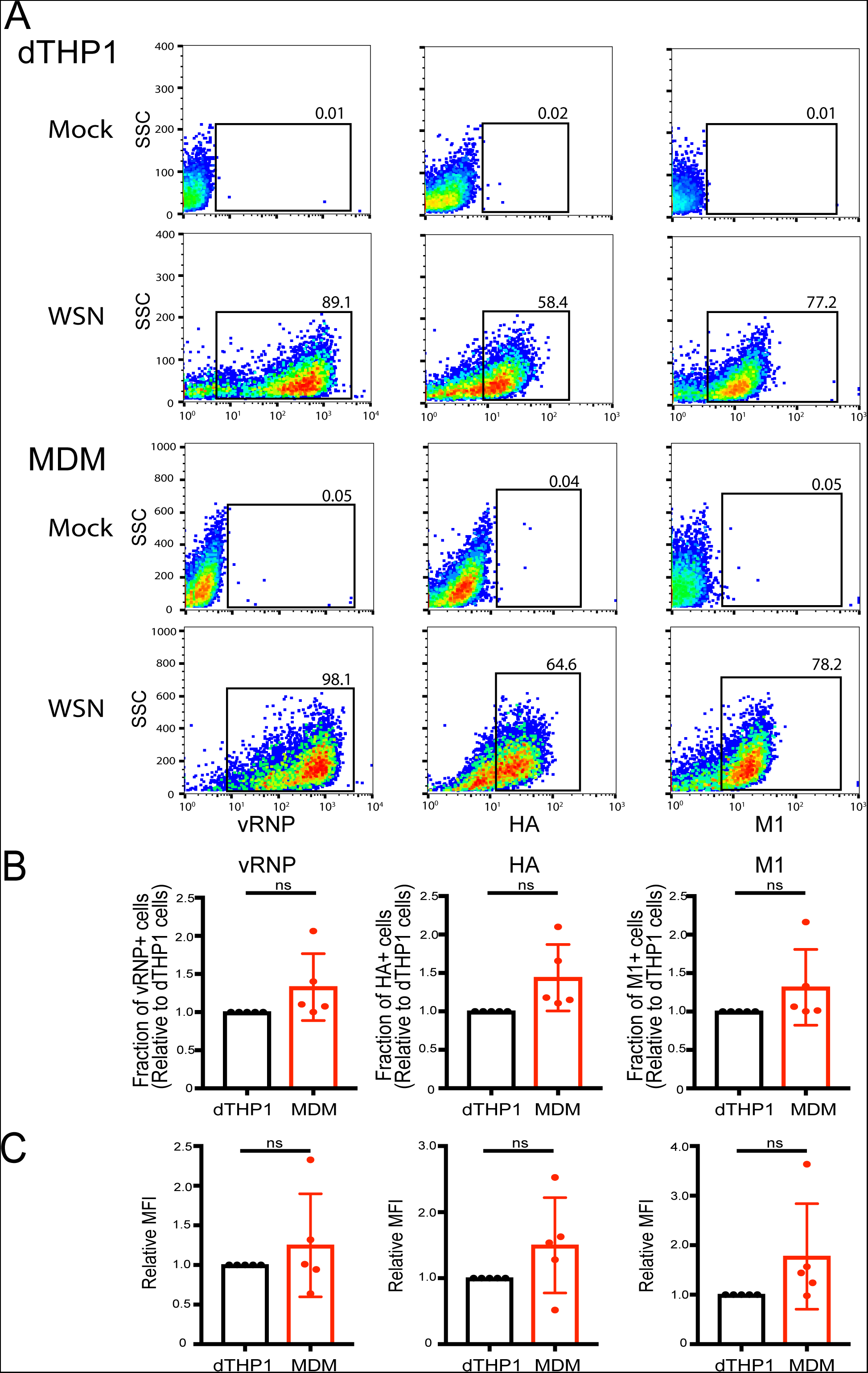
MDM express viral proteins as efficiently as dTHP1 cells. dTHP1 cells and MDM were infected with WSN at MOI 0.01 for 24 hours. Infected cells were fixed, permeabilized and analyzed for cellular levels of vRNP, HA and M1 by flow cytometry. (A) Representative flow plots for mock- and WSN-infected cells are shown. Gates for positive cell populations were set in comparison to mock-infected cells. Due to differences in the side scatter (SSC) profile between dTHP1 cells and MDM, the Y-axis (SSC) range is different between the two cell types. (B) Percentages of cells positive for vRNP, HA, and M1 are compared between dTHP1 and MDM. (C) MFIs (normalized to dTHP1 cells) for all three viral components are shown for positive cell populations (gated in panel A). Data are from at least three independent experiments and are shown as mean +/- SD. ns, non-significant.

**Supplementary Figure 3:**
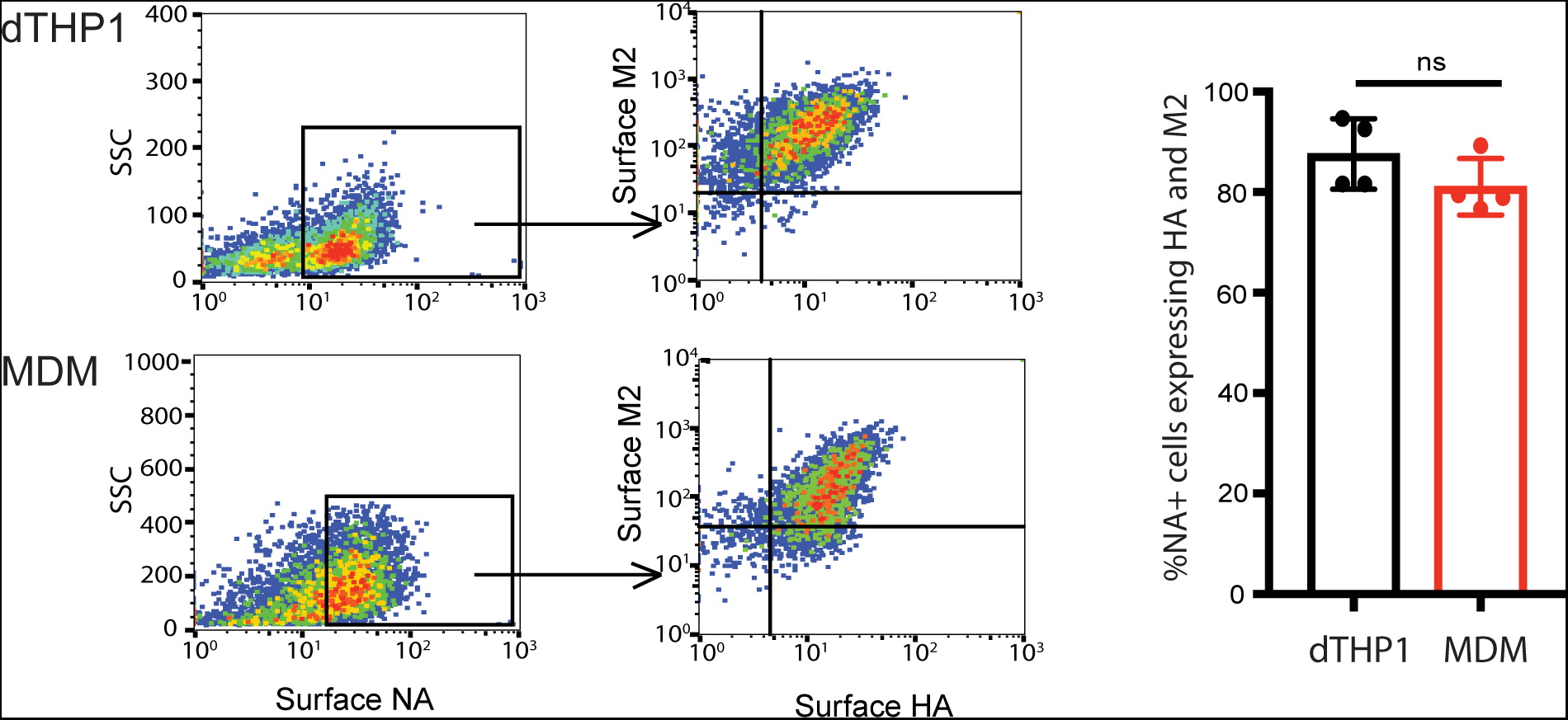
Majority of NA-expressing dTHP1 cells and MDM co-express both HA and M2. dTHP1 cells and MDM were infected with WSN at MOI 0.1 for 16 hours. Cells were fixed and stained for surface HA, M2, and NA. Representative plots are shown in the left panel. % cells expressing HA and M2 within the NA-positive cell population were determined and shown in the right panel. Data are from at least three independent experiments and shown as mean +/- SD. ns, non-significant.

**Supplementary Figure 4:**
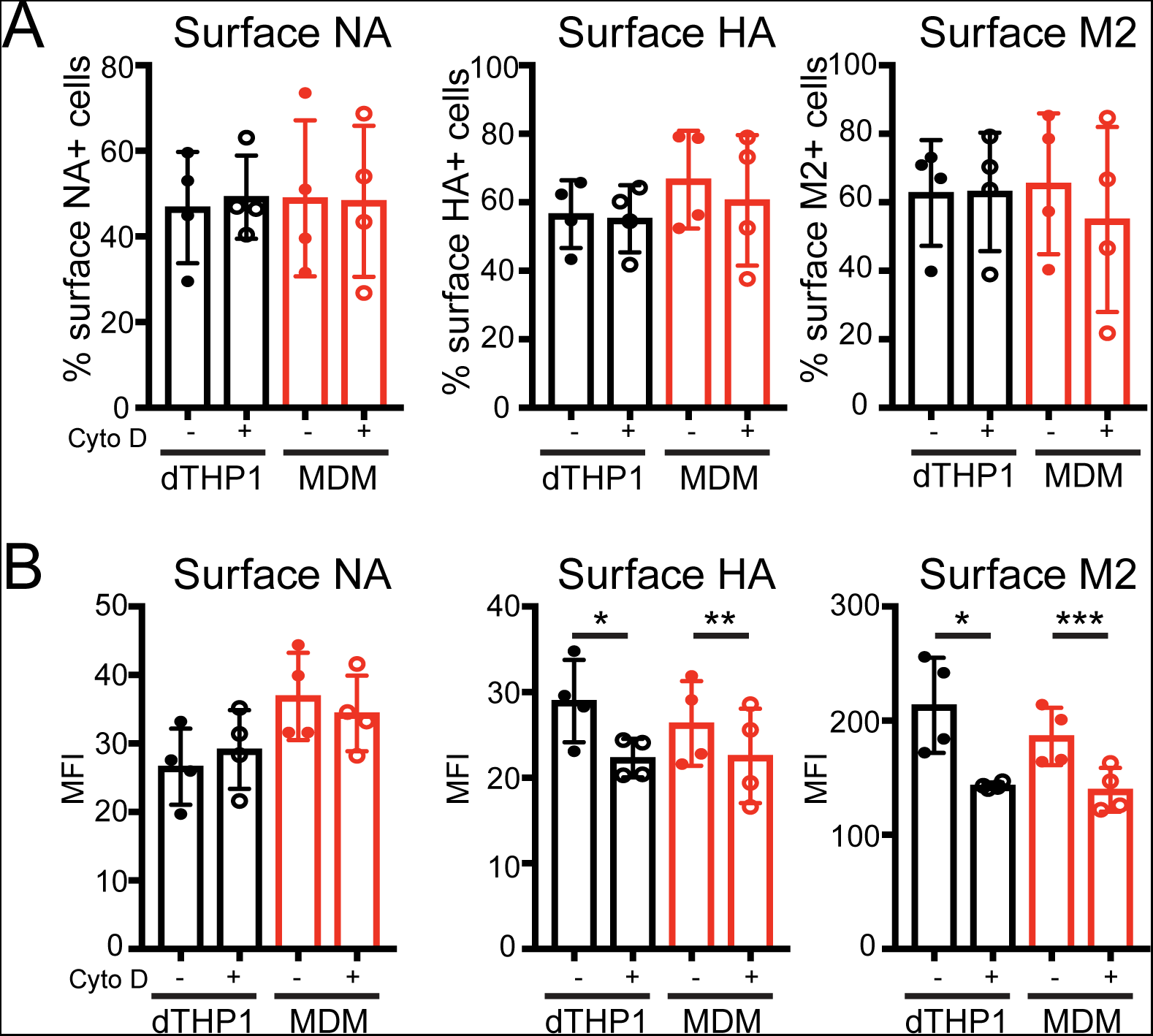
Cytochalasin D treatment does not increase cell surface expression of HA, NA, and M2 in dTHP1 cells and MDM. dTHP1 cells and MDM were infected with WSN at MOI 0.1 for 14 hours. Cells were treated with vehicle control (DMSO) or 20 µM Cyto D for 2 hours. (A) % cells expressing HA, NA, and M2 on the cell surface are shown. (B) MFIs for the indicated proteins in positive cell populations are shown. Data are from three experiments and shown as mean +/- SD. *, p<0.05; **,p<0.01; ***p<0.005.

**Supplementary Methods:** Details of methods for virus infection and detection of infection are described.

## Supplementary Methods

### Virus infection

For virus infection, cells were first washed twice with MEM-BSA medium (1X MEM supplemented with 0.3% BSA (Sigma), 0.23% sodium bicarbonate (Gibco), 1X MEM amino acids (Gibco), and 1X MEM vitamins (Gibco)). The cells were then inoculated with diluted virus at MOI 0.01 or 0.1 for 1 hour at 37°C, washed twice with MEM-BSA, and cultured in MEM-BSA containing 0.2 μg/ml tosylsulfonyl phenylalanyl chloromethyl ketone (TPCK)-treated trypsin (Worthington).

### Infectious virus measurement by plaque assay

Confluent monolayers of MDCK cells (cultured up to passage 40) plated in 24-well plates were washed twice with MEM-BSA and incubated with 100 μl of serial dilutions of virus supernatants for 1 hour at 37°C. Cells were washed once with MEM-BSA and layered with MEM-BSA containing 0.76 ug/ml TPCK-treated trypsin and 1% Seakem GTG agarose /Seaplaque (Lonza). After gelation of agarose, the plates were inverted and incubated at 37°C. After 48-72 hours, the agarose was removed, and the cells were stained with crystal violet (0.1% crystal violet in 20% methanol) for 10 minutes before counting plaques.

### Flow cytometry

For viral protein expression analysis by flow cytometry, virus- or mock-infected cells were detached using 0.06% trypsin-EDTA in phosphatebuffered saline (PBS) and fixed with 4% paraformaldehyde (PFA, Electron Microscopy Sciences) in PBS for 20 minutes at room temperature. After fixation, cells were washed twice with 2% FBS in PBS (FACS buffer). For detecting viral protein expression on the cell surface, cells were subsequently incubated with primary antibodies for 45-60 minutes directly. For staining of intracellular proteins, cells were first permeabilized with 0.1% TritonX-100 in PBS for 5 minutes and then probed with primary antibodies. After washing once with the FACS buffer, cells were incubated with fluorescently labeled secondary antibodies (Invitrogen) for 20-30 minutes. Cells were washed twice with the FACS buffer and analyzed using the FACSCanto flow cytometer (BD Biosciences). Data were analyzed in FlowJo (Treestar), and the positive gates were set using the mock-infected controls.

### Primers for vRNA measurement

The following primers were used for detection of vRNAs:

PB1 forward: 5’-TCAGAGAAAGAGACGAGTGAG-3’, PB1 reverse: 5’-AAACCCCCTTATTTGCATCC-3’

PB2 forward: 5’- GTTGGGAGAAGAGCAACAGC-3’, PB2 reverse: 5’-GATTCGCCCTATTGACGAAA-3’

PA forward: 5’- TGTGCAGCAATGGATGATTT-3’, PA reverse: 5’-TCTCCCATTTGTGTGGTTCA-3’

NP forward: 5’- GCGCCAAGCTAATAATGGTG-3’, NP reverse: 5’-GGAGTGCCAGATCATCATGT-3’

M forward: 5’-GACCRATCCTGTCACCTCTGAC-3’, M reverse: 5’-AGGGCATTYTGGACAAAKCGTCTA-3’

NS forward: 5’- CAGAATGGACCAGGCGATCA-3’, NS reverse: 5’-TAGAGTCTCCAGCCGGTCAA-3’

HA forward: 5’-TAACCTGCTCGAAGACAGAC-3’, HA reverse: 5’-AGAGCCATCCGGTGATGTTA-3’

NA forward: 5’- TTGGTCAGCAAGTGCATGTC-3’, NA reverse: 5’-ACAGCCACTGCTCCATCATC-3’.

